# Neuropathological stage-dependent proteome mapping of the olfactory tract in Alzheime’s disease: From early olfactory-related omics signatures to computational repurposing of drug candidates

**DOI:** 10.1101/2023.10.01.560342

**Authors:** Paz Cartas-Cejudo, Adriana Cortés, Mercedes Lachén-Montes, Elena Anaya-Cubero, Elena Puerta, Maite Solas, Joaquín Fernández-Irigoyen, Enrique Santamaría

## Abstract

Alzheimer’s disease (AD) is the most common form of dementia, characterized by an early olfactory dysfunction, progressive memory loss and behavioral deterioration. Albeit substantial progress has been made in characterizing AD-associated molecular and cellular events, there is an unmet clinical need for new therapies. In this work, olfactory tract proteotyping performed in controls and AD subjects (n=17/group) showed a Braak stage-dependent proteostatic impairment accompanied by the progressive modulation of amyloid precursor protein (APP) and tau functional interactomes. To implement a computational repurposing of drug candidates with capacity to reverse early AD-related olfactory omics signatures, we generated a consensual olfactory omics signatures (OMSs) database compiling differential omics datasets obtained by mass-spectrometry or RNA-sequencing derived from initial AD across the olfactory axis. Using the Connectivity Map (CMAP)-based drug repurposing approach, PKC, EGFR, Aurora kinase, Glycogen synthase kinase and CDK inhibitors were the top pharmacologic classes capable to restore multiple OMSs, whereas compounds with targeted activity to inhibit PI3K, IGF-1, microtubules and PLK represented a family of drugs with detrimental potential to induce olfactory AD-associated gene expression changes. In-vitro validation assays revealed that pretreatment of human neuron-like SH-SY5Y cells with the EGFR inhibitor AG-1478 showed a neuroprotective effect against hydrogen peroxide-induced damage while the pretreatment with the Aurora kinase inhibitor Reversine reduced amyloid-beta (Aβ)-induced neurotoxicity. Taken together, our data pointed out that olfactory omics signatures may be useful as substrates for drug repurposing to propose novel neuroprotective treatments against AD.

**STATEMENTS:** *Data availability statement:* Mass-spectrometry data and search results files were deposited in the Proteome Xchange Consortium via the JPOST partner repository (https://repository.jpostdb.org) with the identifier PXD038061 for ProteomeXchange and JPST001921 for jPOST (for reviewers: https://repository.jpostdb.org/preview/1400199357636bce4231af5 Access key: 8609). The data supporting the findings of this study are available in Supplementary Material. Raw data are available from the corresponding author, upon reasonable request.

*Funding statement:* This work was funded by grants from the Spanish Ministry of Science, Innovation and Universities (Ref. PID2019-110356RB-I00/AEI/10.13039/501100011033) to J.F.-I. and E.S. and the Department of Economic and Business Development from Government of Navarra (Ref. 0011-1411-2023-000028 to E.S.). PC-C was supported by a predoctoral fellowship from the Public University of Navarra (UPNA). ML-M is supported by a postdoctoral fellowship from Miguel Servet Foundation-Navarrabiomed. EA-C is supported by “Programa MRR Investigo 2023” in the framework of the European Union recovery and resilience facility.

*Conflict of interest disclosure:* Authors declare that they have no conflicts of interest/financial disclosures.

*Ethics approval and patient consent statement:* According to the Spanish Law 14/2007 of Biomedical Research, inform written consent from several Spanish Neurological Tissue Banks was obtained for research purposes from relatives of subjects included in this study. According to the Declaration of Helsinki, all assessments, post-mortem evaluations, and experimental procedures were previously approved by the Clinical Ethics Committee of Navarra Health Service (Study code: PI_2019/108).

## INTRODUCTION

Alzheimer’s disease (AD) is the most common form of neurodegenerative dementia in the elderly, coursing with functional and cognitive damage together with a specific neuropathological profile (1). The most relevant pathologic hallmarks of AD are the deposition of extracellular Aβ peptides in plaques and the aggregation of intracellular hyperphosphorylated tau in neurofibrillary tangles (NFTs) (2). Tau deposits spread during disease progression across brain areas in a predictable pattern, enabling an objective neuropathologic classification into different Braak stages (3, 4). Although substantial progress has been made in characterizing AD-related molecular and cellular events, early disease-related processes are not well understood. There is an unmet clinical need to identify not only a combination of new biomarkers for early and presymptomatic diagnostic purpose, but also for assessing novel therapies.

Together with behavioral derangements and memory loss, AD patients present olfactory dysfunction in 90% of the cases (5). This deficit occurs at early stages of the disease, being proposed as premotor sign of neurodegeneration (6, 7). Multiple reports have evidenced neuropathological changes and molecular alterations in olfactory-related areas in human AD brains (8-14). Moreover, the presence and severity of hyperphosphorylated Tau and Aβ in human olfactory bulb (OB) and olfactory tract (OT) reflects the presence and severity of AD in other brain regions (5). Indeed, an axonal degeneration has been detected in the OT from AD subjects (15). It has been shown that the loss of fiber OT integrity correlate with a reduction in glucose metabolism in central olfactory structures as well as a loss of gray matter density in pre-AD mild cognitive impairment (MCI) individuals (16, 17). In addition, early and sequential morphological OT changes correlates with dementia development (18). A relevant approach to map molecular disruptions associated to AD is to generate protein expression profiles during its progression, in well-defined subsequent stages and using early affected brain areas. Proteome-wide exploration based on high-resolution mass-spectrometry is considered an attractive approach to proteotype olfactory areas in different biological conditions (19). Although this unbiased methodology has dramatically increased the biochemical knowledge associated to AD (20, 21), few studies have deeply analyzed the progressive proteostatic imbalance of early-affected areas with the aim to unveil incipient neurodegenerative changes in AD phenotypes (11, 14, 22). We consider that the molecular characterization of the OT progressive neurodegeneration that occurs in a Braak stage-dependent manner, will complement the integrated view of the biochemical pathways affected during the olfactory pathophysiology of AD, building the foundations towards the implementation of computational repurposing of drug candidates with capacity to reverse AD-related olfactory omics signatures. In this work, we combine neuropathological diagnosis, quantitative proteomics, physical/functional interaction data and large-scale perturbation databases to understand how the molecular pathways harboured in the OT are chronologically regulated during AD progression as well as to perform drug repurposing and side-effect prediction for potential new treatments.

## METHODS

### Human samples

According to the Spanish Law 14/2007 of Biomedical Research, inform written consent from several Spanish Neurological Tissue Banks (Hospital Clinic-Institut d’Investigacions Biomédiques August Pi i Sunyer-IDIBAPS, IDIBELL Biobank and the Biobank from Navarrabiomed) was obtained for research purposes from relatives of subjects included in this study. According to the Declaration of Helsinki, all assessments, post-mortem evaluations, and experimental procedures were previously approved by the Clinical Ethics Committee of Navarra Health Service (Study code: PI_2019/108). Post-mortem brain processing, extraction and fixation were performed as previously described (23). The neuropathological diagnosis was carried out according to the current neuropathological guidelines. Cases with associated proteinopathies (i.e. TDP-43 pathology) were not included in this study. Controls did not suffered from neurological/mental diseases, and the neuropathological exploration revealed no alterations excepting for small blood vessel disease in some cases. Thirty-four human OT samples from post-mortem subjects were employed for proteomics screening of which 17 of them were controls (n=6F/11M; mean age ± SD 61.7 ± 10.5 years) and 17 AD (n=4F/13M; mean age ± SD 74.4 ± 7.8 years) (Table 1).

**Table 1.** Description of olfactory tract samples included in this study. PMI; post-mortem interval.

|  | Case | Sex | Age | PMI | Neuropathological diagnosis |
| --- | --- | --- | --- | --- | --- |
| **Controls** | A1423 | W | 54 | 8h | non-AD |
|  | A1218 | W | 65 | 3h 45m | non-AD |
|  | A1403 | W | 51 | 4h | non-AD |
|  | A1576 | W | 60 | 12h | non-AD |
|  | A9/43 | W | 51 | 5h 30m | non-AD |
|  | A15/7 | W | 54 | 13h 40m | non-AD |
|  | C048 | M | 68 | 10h | non-AD |
|  | C064 | M | 87 | 21h | non-AD |
|  | A1404 | M | 65 | 9h 35m | non-AD |
|  | A1419 | M | 56 | 8h 50m | non-AD |
|  | A1412 | M | 77 | 6h 10m | non-AD |
|  | BCN473 | M | 65 | 3h 05m | non-AD |
|  | A1277 | M | 74 | 9h 25m | non-AD |
|  | A1368 | M | 45 | 18h 30m | non-AD |
|  | A1438 | M | 67 | 5h 50m | non-AD |
|  | A1377 | M | 59 | 8h 30m | non-AD |
|  | A15/30 | M | 51 | 10h | non-AD |
| **AD** | EA1 | W | 73 | 3h 30m | Braak stage VI |
|  | EA6 | W | 77 | 6h | Braak stage VI |
|  | A1429 | W | 78 | 3h 15m | Braak stage V |
|  | A1242 | W | 82 | 17h 15m | Braak stage IV / B |
|  | EA2 | M | 75 | 4h | Braak stage VI / C |
|  | EA4 | M | 75 | 4h 15m | Braak stage VI / C |
|  | EA5 | M | 84 | 5h | Braak stage VI / C |
|  | EA7 | M | 80 | 6h 30m | Braak stage V / C |
|  | EA10 | M | 86 | 9h | Braak stage VI / C |
|  | A1247 | M | 81 | 5h 5m | Braak stage III / A |
|  | A1517 | M | 84 | 20h 45m | Braak stage III / A |
|  | A9/92 | M | 81 | 7h 30m | Braak stage III / A |
|  | A9/95 | M | 63 | 3h 50m | Braak stage I / A |
|  | A9/121 | M | 63 | 11h 30m | Braak stage V / B |
|  | A11/13 | M | 70 | 2h | Braak stage I / A |
|  | A11/75 | M | 61 | 4h 30m | Braak stage I |
|  | A11/85 | M | 59 | 4h 15m | Braak stage I |

### OT Proteome-wide mapping

Protein extraction, sequential window acquisition of all theoretical fragment ion spectra mass spectrometry (SWATH-MS), MS/MS library generation, proteome quantitation and data analysis was performed as previously described (23). Mass-spectrometry data and search results files were deposited in the Proteome Xchange Consortium via the JPOST partner repository (https://repository.jpostdb.org) with the identifier PXD038061 for ProteomeXchange and JPST001921 for jPOST (for reviewers: https://repository.jpostdb.org/preview/1400199357636bce4231af5 Access key: 8609).

### Bioinformatics and statistical analysis

The quantitative data obtained by PeakView® were analyzed using Perseus software (version 1.6.15.0) (24) was used to perform statistical analysis and visualization of the obtained data. Welch-s t-test was used for direct comparisons between controls, and AD staging. Statistical significance was set at p-value lower than 0.05 in all cases and 1% peptide FDR threshold was considered. In addition, proteins were considered significantly differentially expressed when their absolute fold change was below 0.77 (down-regulated proteins) and above 1.3 (up-regulated proteins) in linear scale. Boxplots were performed with R software (v 4.1.2). The association of the differentially expressed proteins with specifically dysregulated regulatory/metabolic networks in OT human samples was analyzed using QIAGEN’s Ingenuity Pathway Analysis (IPA; QIAGEN Redwood City). This software calculates significance values (p-values) between each biological or molecular event and the imported molecules based on the Fisher’s exact test (p ≤ 0.05). Metascape (25) was also used to extract biological information associated to proteome functionality using default settings (minimum overlap: 3; minimum enrichment: 1.5; P < 0.01). As a drug repurposing approach, we interrogated the Connectivity Map (CMap) dataset that contains the effects of over 5000 small molecules on the transcriptome of human cell lines and looked for molecules which effects on transcription mimic or oppose those of the olfactory omics signatures (OMSs). Sixteen independent OMSs were used as inputs to query CMap using its clue.io tool (https://clue.io), in order to obtain similarity scores for the signatures associated to all the perturbagens (compounds, CMap classes and knock-down of genes) available at CMap (26, 27). The Query CMap tool was used, employing Gene expression (L1000), Touchstone and individual query as query parameters. Similarity scores were generated, and results sorted based on their scores and the type of perturbagen.

### SH-SY5Y Neuroblastoma Cell Culture

The human Neuroblastoma clonal SH-SY5Y cells were obtained from American Type Culture Collection (ATCC) (CRL-2266). Cells were cultured in DMEM/F12 (Gibco, ref. 10565018) containing GlutaMAXTM and was supplemented with 10% FBS (Gibco), penicillin (100 U/mL, Gibco) and streptomycin (100 U/mL, Gibco). Cells were maintained at 37°C in saturated humidity (5% CO2). Non-differentiated cells were plated on 48-well plates (Corning), at a density of 5 × 104 cells per well and were used 24 h after seeding.

### Aβ_25–35_ Soluble Species Preparation

Aβ25–35 fragment was purchased from Sigma-Aldrich laboratories (ref. A4559-1MG). 1 mg of Aβ*_25–35_* fragment was dissolved in 1 mL of type II water and was frozen at -20°C until further use. In order to obtain toxic oligomeric forms, Aβ*_25–35_* at a concentration of 10 μM was incubated in PBS at 37°C for 2 hours prior cell treatment.

### Cell Treatment and MTT Cell Viability Assay

Compounds including AG-1478 (HY-13524), reversine (HY-14711), indirubin (HY-N0117) and GSK-3β inhibitor II (HY-130795) were acquired from MedChemExpress (New Jersey, USA) in a 1 mL, 10 mM in DMSO format. SH-SY5Y cells were treated 30 minutes with each tested drug at a different concentration: AG-1478 (50, 100, 200, 400, 800, 1000 nM), Reversine (50, 100, 200, 400, 800, 1000 nM), Indirubin (5, 10, 20, 40, 80 and 100 nM) and GSK-3β inhibitor II (0.5, 1, 5, 10, 20 and 50 nM). Then, hydrogen peroxide (80 μM) or Aβ*_25–35_* (10 μM) was added for 24 hours. Cell viability was examined by the 3,4,5-dimethylthiazol-2-yl-2,5-diphenyltetrazolium bromide (MTT) assay. MTT assay is a colorimetric assay for measuring the activity of cellular enzymes in living cells in response to potential toxic. After the cell treatment, culture medium was replaced by a solution of 0.5 mg/mL MTT (Sigma-Aldrich) in DMEM (Gibco, 31053-028). Cells were incubated with MTT solution for 2 h in the cell incubator (5% CO2 and 37°C). Then, the MTT solution was discarded and DMSO was added to the wells. Aliquots were transferred to a 96-well plate, and absorbance was measured at 590 nm in a plate reader. Results were expressed as percentages of non-treated control cells.

## RESULTS

### Olfactory tract proteostatic imbalance during AD progression

To characterize and determine the complexity and progression of neuropathological-associated changes, the OT site-specific proteomic fingerprint was monitored across Braak staging using SWATH-MS quantitative proteomics. Among 1835 quantified proteins across all experimental groups, 201 proteins tend to be differentially expressed between controls and AD phenotypes (Supplementary figure 1 and Supplementary table 1). However, the stage-dependent analysis revealed that 116, 144, and 335 olfactory proteins were differentially expressed (DEPs) in Braak I-II, Braak III-IV, and Braak V-VI stages respectively (Figure 1A). The differential expression distribution between downregulated and upregulated proteins was very similar across Braak I-IV grading (around 35% downregulated proteins in Braak I-II and III-IV stages). This percentage reached 45% in Braak V-VI stage (Figure 1A, Supplementary Table 1). Clustering analysis revealed protein subsets with specific protein expression profiles across neuropathological grading (Figure 1B). Specifically, we observed protein clusters with a significant aberrant expression in Braak I-II or in Braak III-IV that followed different trajectories across late stages (Figure 1C and D) as well as a pair of clusters composed by proteins exclusively modulated in Braak V-VI stages (Figure 1E). Interestingly, the overlap across all stages was very low (Figure 1F). Only two OT DEPs overlapped between all stages (*ALDH16A1*, *COL6A1*) whereas 7 proteins were deregulated across Braak I-IV stages (*COPB2*, *PSMB4*, *PPP1R1B*, *AKR7A3*, *RPL7*, *NID2*, *TINAGL1*) (Supplementary Table 1), highlighting the molecular heterogeneity in AD progression at the level of this olfactory structure. However, part of the OT DEPs identified in different Braak stages were involved in the same biological processes (Figure 1G), evidencing a more extensive functional overlap across neuropathological stages.

**Figure 1.**
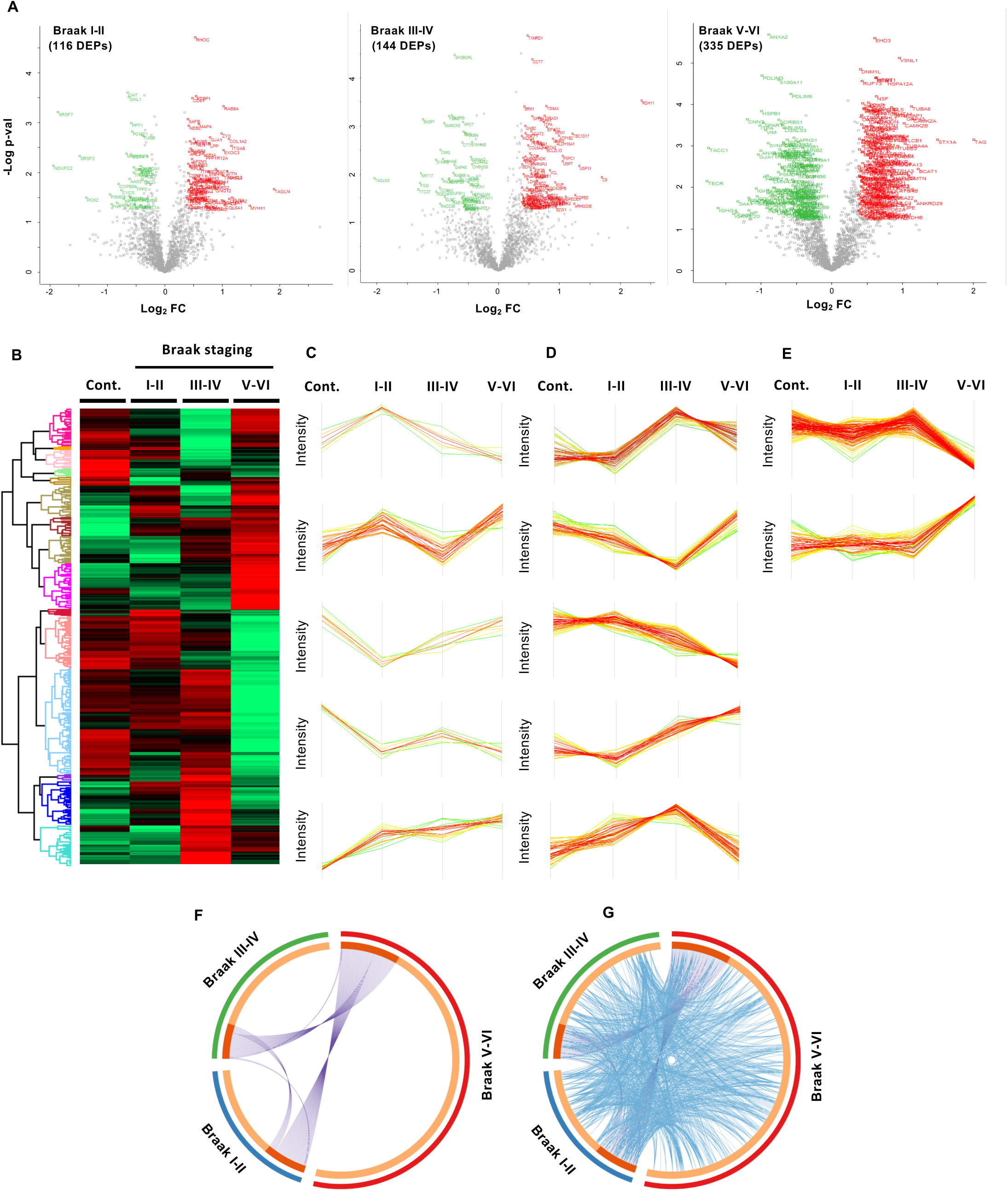
Neuropathological-stage dependent proteomic analysis of the OT in AD. A) Volcano-plots representing the progressive proteostatic imbalance across Braak stages. B) Heatmap representing the diferential OT proteotyping in AD across Braak stages. C) Protein clusters modulated in AD subjects (Braak I-II). D) Protein clusters modulated in AD subjects (Braak III-IV). E) Protein clusters modulated in AD subjects (Braak V-VI). F) Circos-plot representing the OT deregulated proteome shared across neuropathological stages (purple lines). G) Circos-plot representing the functional overlap (blue lines) between dysregulated proteomes across Braak stages at the level of the OT.

### Functional commonalities and differences across Braak stages in the olfactory tract

To decipher specificities and singularities associated to initial, intermediate, or advanced Braak stages at the level of the OT, stage-dependent proteomic datasets were functionally analyzed. As shown in Figure 2A, common pathways and processes were imbalanced across all Braak stages. Specifically, lipid metabolism, aminoacid metabolism, vesicle-mediated transport and signaling by receptor tyrosine kinases and Rho GTPases between others were commonly altered during AD olfactory neurodegeneration (Figure 2A and Supplementary Table 2). Process such as signaling by ROBO receptors, complement cascade, mRNA 3’-end processing, RAS regulation by GAPs and MAPK6/MAPK4 signaling were specifically altered in initial Braak I-II stages (Figure 2A and Supplementary Table 2). However, alterations associated to signaling by PDGF, signaling by ALK, platelet homeostasis and tRNA aminoacylation were observed in intermediate Braak III-IV stages (Figure 2A and Supplementary Table 2). A widespread metabolic and postransductionally molecular events (glycosylation, phosphorylation) were exclusively detected in advanced Braak V-VI stages (Figure 2A). At subcellular level, common components were differentially affected across AD staging (Figure 2B). As expected, due to OT is mainly composed by axonal bundles derived from the OB, GO terms related to cytoskeleton such as cell-cell junction, basement membrane, actin complex, focal adhesion, axon, and cell leading edge were significantly enriched from initial to advanced stages (Figure 2B). However, cytosolic ribosome, centrosome and Golgi-associated vesicles were specifically enriched in Braak I-II stage while GO terms related to spindle midzone, spliceosomal complex and lipid droplets were deregulated in Braak III-IV stages (Figure 2B). Again, multiple alterations were specifically mapped in Braak V-VI, widely affecting subcellular compartments involved in glial cell projection, mitochondrial respiratory chain complexes, perineuronal net, dendritic shaft, clathrin-sculpted glutamate transport vesicle, myelin sheath, axonal growth cone and presynaptic endocytic zone between others (Figure 2B). According to AlzData (http://www.alzdata.org/) (28), part of the differential OT proteome observed in our AD cohort has been previously linked to AD (Supplementary Table 3). Focusing on Braak I-II stages, the expression of protein-coding genes corresponding to 40 and 27 DEPs correlates with AD pathology in Aβ and/or Tau mouse models respectively (Figure 3A and Supplementary Table 3). Moreover, the protein coding genes for 44 DEPs (38% of the differential dataset) are also differentially expressed in AD mouse models before AD pathology is established (Figure 3A and Supplementary Table 3), suggesting an early role during the neurodegenerative process.

**Figure 2.**
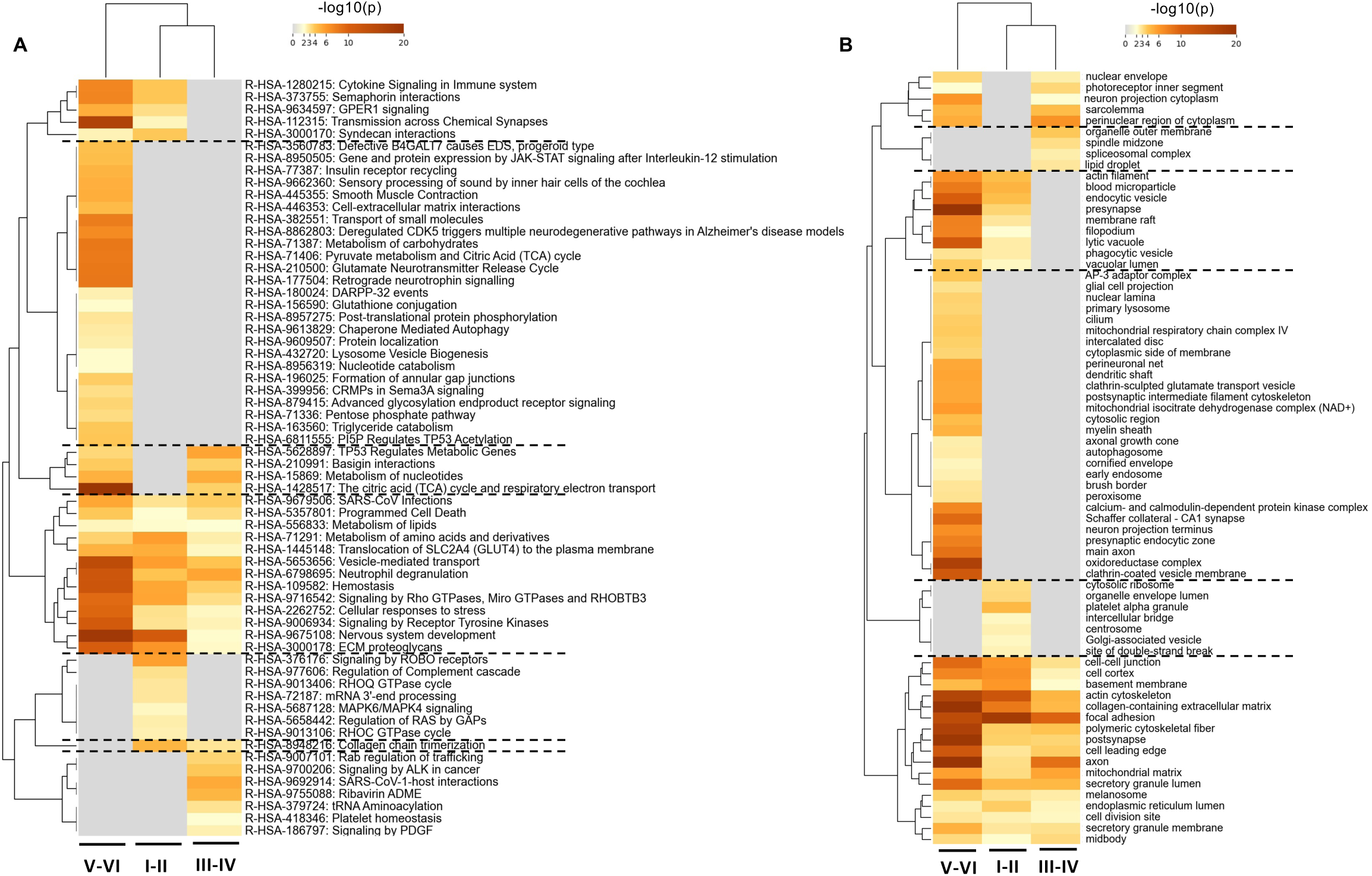
Functional impact of the deregulated OT proteostasis across AD Braak stages. A) Functional mapping of disrupted OT proteome across AD neuropathological staging at pathway level. B) Functional mapping of disrupted OT proteome across AD grading at subcellular level.

**Figure 3.**
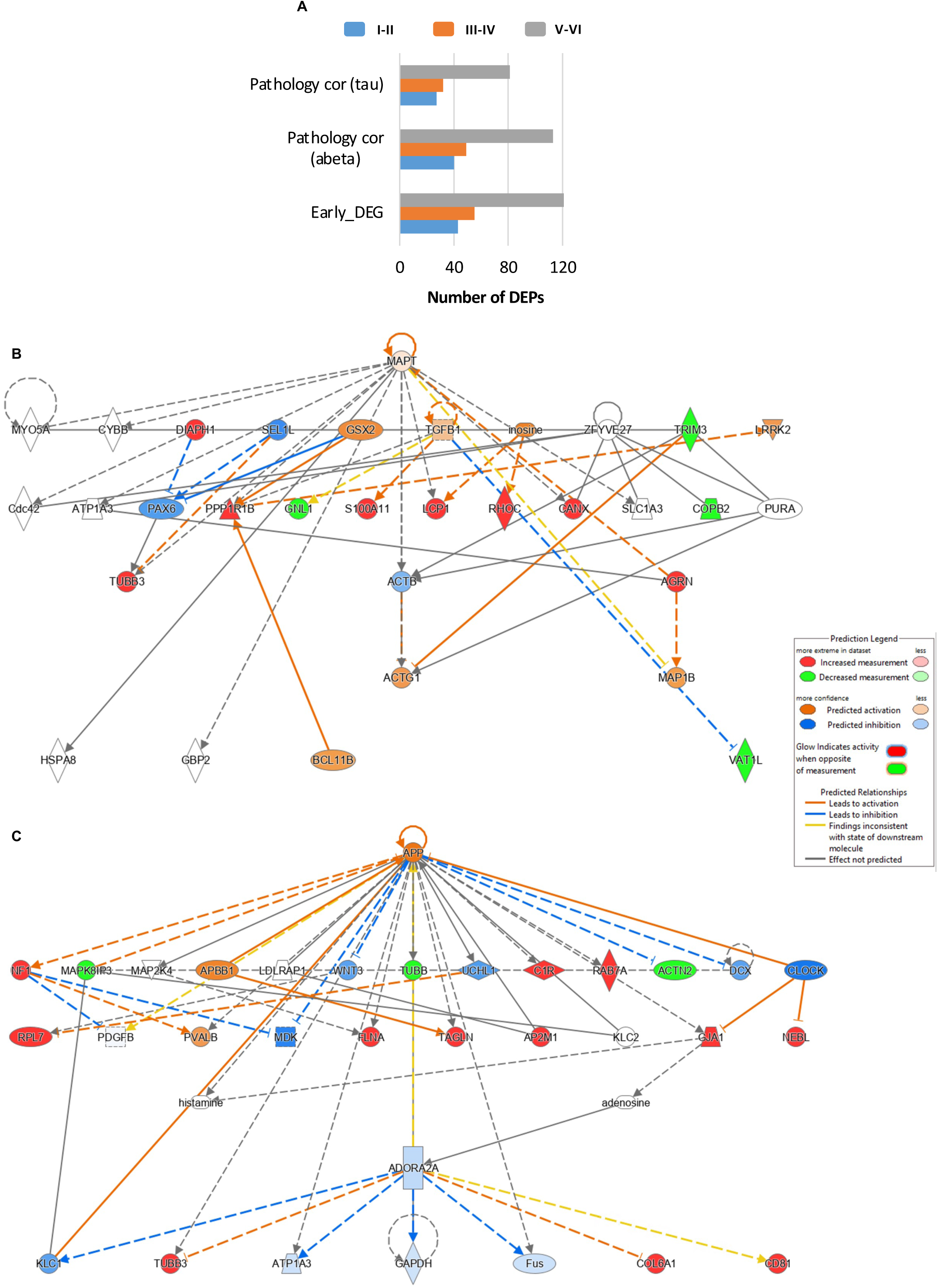
Interlocking of stage-dependent deregulated OT proteomes with Aβ and Tau biology. A) Integrative analysis of our OT proteome data derived from AD subjects with AlzData database (http://www.alzdata.org/). B) Deregulation of *MAPT* functional network in initial Braak I-II stages at the OT level according to IPA knowledgebase. C) Deregulation of OT *APP* functional network in initial Braak I-II stages according to IPA knowledgebase. DEGs; differentially expressed genes.

### Alterations in AD-related protein interactomes during OT neurodegeneration

We wanted to know whether deregulated OT protein subsets were directly associated to neuropathological interactomes. A systems-biology approach allowed us to partially characterize the modulation of functional protein networks associated to APP and MAPT (tau) (Figure 3B and C). In addition, according to BIOGRID experimental repository (29), part of the OT deregulated proteome physically interacts with APP and Tau proteins (Supplementary figure 2 and 3). Based on previous high-quality proteomic maps (30-32), some DEPs were also components of human Aβ plaques (Figure 4A and Supplementary Figures 4, 5 and 6) or neurofibrillary tangles (NFTs) (Supplementary figure 7). As an example, COL6A1, an overexpressed protein across all AD stages, has been previously detected in Aβ plaques (Figure 4A). MAP4, SRSF7 (deregulated OT proteins in Braak I-II stages) and OXCT1 (down-regulated in initial and advanced stages) are physical interactors of phospho-Tau in NFTs (Figure 4B).

**Figure 4.**
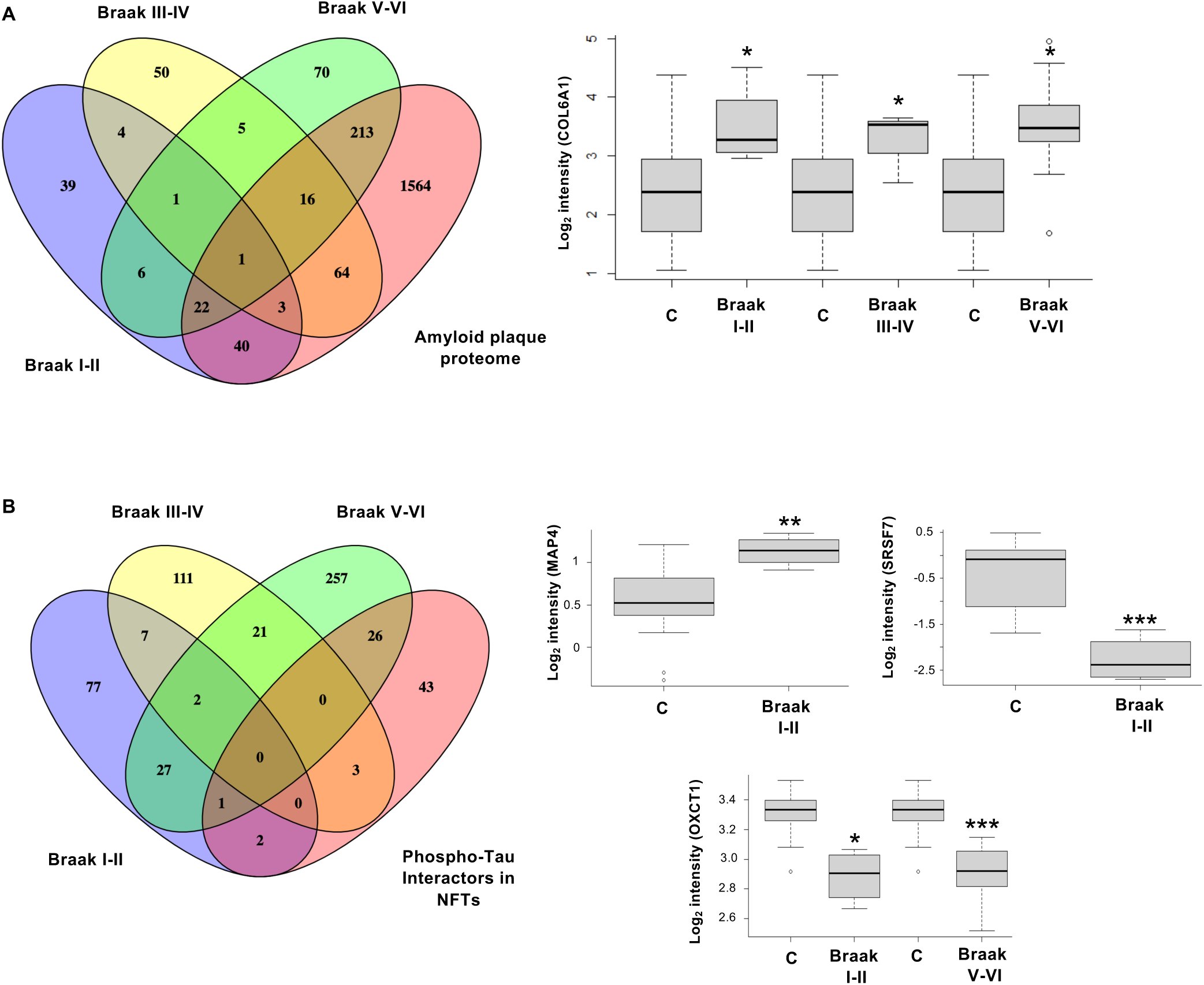
OT Differential proteins as components of AD plaques and NFTs. Venn diagram showing the overlap between OT altered proteins and protein components of Aβ plaques (A) and pTau interactors in NFTs (B).

### Functional commonalities and differences across initial AD grading: A neuroanatomical proteomic integration across different brain areas

To establish a functional relationship between the OT and other olfactory AD-affected regions at the proteome level, a traceability analysis was done comparing the OT differential protein set with respect to altered proteins previously detected in functionally related structures such as the olfactory bulb, entorhinal cortex, and hippocampus derived from subjects with initial AD (Braak stages I-II (11, 14, 22). Besides the functional overlap observed across primary and secondary olfactory areas (Supplementary Figure 8a), a minimal protein set was commonly deregulated between different structures (Supplementary Figure 8b) suggesting the absence of conserved transcriptional programs that may be activated/deactivated across structures during olfactory neurodegeneration in initial AD stages.

### Drug prediction using AD-related olfactory omics signatures (OMSs)

Databases containing transcriptional signatures of human cell lines subjected to a specific perturbation (such as drugs or genetic manipulations) have become an ideal platform for data mining in neuroscience research. Specifically, the Connectivity Map (CMap) is a chemogenomic-based drug repurposing tool that stores over 1.5 M signatures derived from a wide range of human cancer cell lines exposed to around 5000 compounds and 3000 genetic perturbations (26, 27). In our case, we used an olfactory proteotranscriptomic-based computational drug-repurposing method to identify drugs that induce a gene expression signature which negatively correlates with early AD phenotype as a strategy for unveiling potential new therapies. First, we compiled the current OT proteomic study as well as other proteomic or transcriptomic studies performed in different primary and secondary olfactory areas (olfactory mucosa, olfactory bulb, entorhinal cortex, hippocampus) in initial stages of AD (11, 14, 22, 33, 34). The OT (Braak I-II) differential dataset was also divided in different lists depending on ALZData parameters (Aβ or Tau correlation in AD murine models, early differentially expressed genes) and on the presence of DEPs in Aβ-plaques, according to previous proteomic evidence obtained by laser-capture microdissection approaches in human AD (30, 31). In total, our consensual olfactory omics signatures (OMSs) database included 16 differential datasets derived from Braak I-II, MCI or AD mild dementia (CDR 1) (Supplementary Table 4). Based on the CMap framework, the connectivity score (CS) is a standardized statistic parameter ranging from −100 to 100. A highly negative score indicates that the drug-gene signatures are opposing in which genes that are upregulated in AD phenotypes are downregulated by the compound and vice versa. Said that, we queried the Cmap to find: i) drugs that potentially reverse OMSs associated to initial AD (beneficial effect prediction; CS lower than -90), ii) drugs that potentially mimic the olfactory neurodegeneration observed in AD (Side-effect prediction; CS higher than +90) and iii) specific genes whose sequence-specific gene silencing tend to reverse AD expression signature under knock-down conditions (potential therapeutic targets; CS lower than -90) (Figure 5A). Some drug pharmacological classes and potential therapeutic genes were significantly represented across OMSs with a lowest CS, indicating that this type of interventions could be good candidates to effectively reverse part of initial AD features at olfactory level (Supplementary Tables 5-7). PKC, EGFR, Aurora kinase, Glycogen synthase kinase and CDK inhibitors were the top pharmacologic classes capable to restore multiple OMSs (Figure 5B and C). It is important to note that compounds with targeted activity to inhibit PI3K, IGF-1, microtubules and PLK appeared with CS higher than +90, representing a family of drugs with detrimental potential to induce AD-associated gene expression changes (Figure 5D and E). Interestingly, expression silencing approaches of genes poorly associated to AD mechanisms (Figure 5F) may be considered as potential therapeutic targets, based on their ability to reverse olfactory molecular profiles in early AD. 6 out of 9 genes (Figure 5F) are expressed in multiple regions in human brain (35). We wanted to assess how drugs that have been obtained by our repurposing approach across all OMSs were prioritized by clinical trials for AD or associated symptoms. For that, we downloaded all available clinical trial information for AD, MCI and dementia from the Clinicaltrials.gov website, resulting in 3079, 1260 and 4436 records respectively. The compilation of data related to drug/compound interventions leaved 1534 records (search done in March, 2023). In our case, 556 drugs/compounds presented a CS lower than -90 in at least one olfactory dataset. The intersection of the clinical trial drug list and the OMSs-derived repurposing candidate list indicated an overlap of three compounds: Digoxin (derived from Aβ Corr and HC Braak I-II datasets and used in NCT00563732 and NCT00831506 trials), Equol (derived from Aβ Corr, Tau Corr and Early DEGs datasets and used in NCT03101085 and NCT02142777 trials) and estradiol (derived from Aβ Corr, Tau Corr and Early DEGs datasets and used in NCT03681691 and NCT00066157 trials). We decided to focus our attention in the most represented candidates across OMSs that belong to top pharmaceutic classes derived from our analysis: Tyrphostin AG-1478 (EGFR inhibitor), Reversine (Aurora kinase inhibitor), Indirubin (CDK inhibitor) and GSK-3β inhibitor II (PKC inhibitor) (Figure 6A). MTT assay was performed in order to determine the range of concentrations that does not lead to cell death in human neuroblastoma cells (SH-SY5Y). Indirubin showed a marked neurotoxicity in all concentration assayed, therefore, this compound was discarded from the study (Supplementary figure 9). The rest of the compounds did not show any toxicity at the concentrations that were chosen for further studies (Supplementary figure 9). We used hydrogen peroxide and Aβ-induced neurotoxicity, with the aim to verify the potential neuroprotective effect of our compound list against AD pathology. While GSK-3 inhibitor II did not show any protective capacity (data not shown), AG-1478 at the highest concentration increased cell viability upon hydrogen peroxide treatment and Reversine showed neuroprotection upon Aβ insult (Figure 6B).

**Figure 5.**
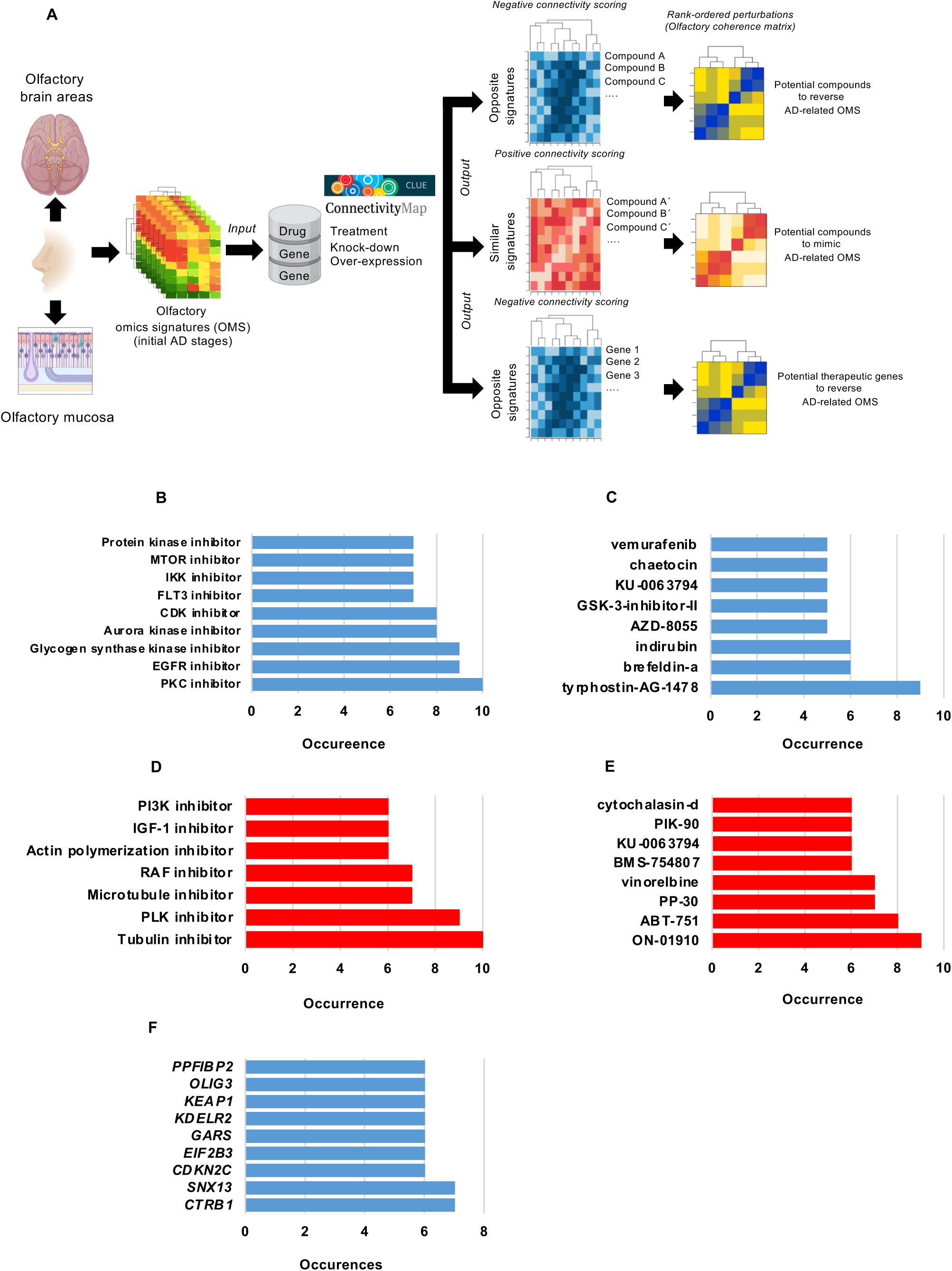
Olfactory omics signatures (OMS) as a template for drug repurposing in AD. A) Drug repurposing workflow used in this study. B) Top pharmacological classes with potential capacity to reverse part of our OMSs database. C) Part of the compounds with predictive potential to reverse OMSs. D) Top pharmacological classes with potential to mimic olfactory AD omics signatures. E) Specific compounds with capacity to induce olfactory AD omics signatures. F) Genes considered as potential therapeutic targets, based on their predictive potential ability to reverse olfactory molecular profiles in early AD when its expression is inhibited. Occurrence refer to the number of OMSs targeted by each pharmacological class or compound in each case.

**Figure 6.**
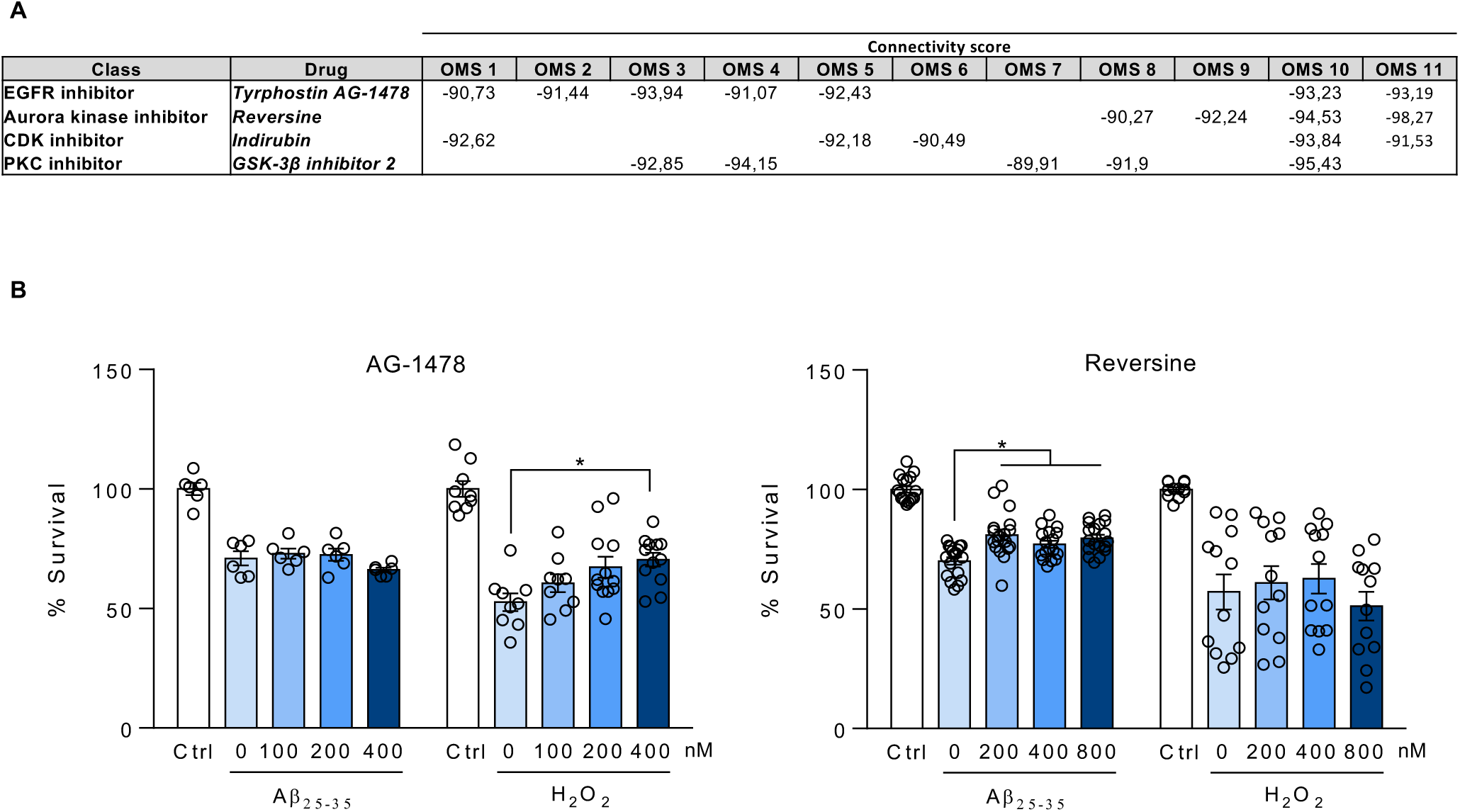
Candidate selection and in vitro validation assays. A) Table showing the four pharmacological classes selected from our workflow. The connectivity score (between -90 and - 100) is indicated for each OMSs: OMS1 (Aβ corr OT), OMS2 (Aβ or Tau corr OT), OMS3 (early DEGs), OMS4 (OB and OT AD I-II), OMS5 (OT AD I-II), OMS6 (Tau Corr OT), OMS7 (OB AD I-II), OMS8 (olfactory mucosa AD), OMS9 (fibro/stromal cells OM AD), OMS10 (GBC cells OM AD), OMS11 (myofibroblasts OM AD). B) In vitro experiments showing the effect of pretreatment of SH-SY5Y cells with AG-1478 (left panel) and Reversine (right panel) upon hydrogen peroxide and Aβ insults. Data was analyzed using One-way ANOVA. * indicates p<0.05.

## DISCUSSION

Based on the involvement of the olfactory dysfunction in AD, the deployment of quantitative olfactory proteomics across AD staging may help to understand the early smell impairment that occurs in this neurodegenerative disease. Multiple neuroproteomic studies have been performed in brain areas differentially affected by the disease (36). However, the molecular knowledge generated by high-throughput approaches associated to the olfactory axis in AD is limited (11, 37). One of the goals of the present study was to generate comprehensive functional data on protein subsets associated to the progression of the neurodegenerative process across the OT region. We have observed that the chronological molecular perturbation in the OT is clearly different to the proteome modulation across initial Braak stages at the level of OB, entorhinal cortex or hippocampus (11, 14, 22), suggesting the absence of coordinated transcriptional programs that may be modulated across brain structures during olfactory neurodegeneration in initial AD stages. Albeit multiple features in OT proteostatic metabolism have been unveiled, we considered that several limitations warrant discussion. It is important to note that low-abundant proteome as well as hydrophobic proteins and receptors that may be relevant during the olfactory neurodegenerative process are not extensively represented in our dataset due to technological issues. In addition, the OT proteomic profiles are limited to bulk protein abundance, avoiding the specific-cell type mapping of molecular changes. However, our study has identified a novel resource of potential biomarkers as well as potential targets that may constitute the basis of innovative intranasal treatments against AD (38). In our case, we have applied a drug repurposing approach through the Cmap framework in which molecular expression changes following a genetic or pharmacologic perturbagen in human cell lines may be associated with AD-associated expression changes to identify unexplored compounds with capacity to partially normalize gene expression. We are aware that Cmap is constituted by transcriptional fingerprints derived from human cancer cell lines, so we cannot exclude that reversion of omics profiles may not be perfectly adapted to primary and secondary olfactory areas. Said that, gene expression related to neurotransmission are not widely represented in our workflow. Current FDA-approved drugs such as acetylcholinesterase (AChE) inhibitors, receptor antagonist or anti-Aβ passive immunizations are mainly focused on neurotransmission modulation, obtaining partially increasing cognitive functions but failing in the prevention of AD progression (38, 39). However, downstream events associated to apoptosis, proliferation, differentiation and inflammation are highly represented in Cmap, opening the door to novel combinations that might trigger the multi-level downstream signals beyond the synaptic plasticity. In fact, similar workflows employing transcriptomic profiles (40-42), gene coexpression networks (43), molecular network modeling (44) and protein-protein interaction networks (45) have been used to discover novel drug candidates for AD as well as for other neurological syndromes (46, 47). Nevertheless, few studies developed further in vitro and/or in vivo validation for computational drug repurposing predictions (42, 44). Our rationale was based on the discovery of compounds/small molecules with capacity to restore the initial imbalance in olfactory metabolism leading to block the progressive neuropathological alterations that accompany the cognitive impairment. For that, we compiled 16 OMSs derived from different areas from the olfactory axis, covering olfactory mucosa, OB, OT, entorhinal cortex and hippocampus in order to identify and prioritize drug compounds for partially relieving AD. Using this workflow, the drug prediction panel from Cmap framework was totally independent of drug known functions and uniquely based on an unsupervised analysis of AD-related OMSs and drug-related differential gene expression. We identified multiple drugs with reversion capacity of pathological olfactory omics signatures. Three of them are currently used in clinical trials against AD, partially validating our olfactory proteotranscriptomic-based computational drug-repurposing. We established in vitro validation for four top-hits with different kinase inhibition capacity (PKC, EGFR, Aurora kinase and CDK inhibitor). In line with our findings, the pharmacological action of PKC/GSK3 inhibitors has been also previously proposed in drug repurposing for AD (40). Tyrphostin AG-1478 (EGFR inhibitor) and Reversine (Aurora kinase inhibitor) showed protective signals in our in-vitro setup. Interestingly, repurposing of EGFR inhibitors in AD has been recently postulated (48). Multiple studies have demonstrated beneficial effects of EGFR inhibitors in AD models based on their behavioral/cognitive enhancing, anti-neuroinflammatory and anti-amyloidogenic activities, autophagy enhancement effect, and anti-astrogliosis (49). AG-1478 blocks the stimulation of release of large soluble derivatives of APP by EGF in epidermal cells (50). Our experimental results showed that AG-1478 pretreatment was not able to restore the survival potential of SH-SY5Y cells upon Aβ_25-35_ stimulation. However, AG-1478 counteracted the oxidant capacity of the hydrogen peroxide-induced damage, significantly increasing the survival potential. In accordance with these data, AG-1478 administration has been shown to ameliorate ROS accumulation and ER stress in diabetic mice (51). The antioxidant role together with its ability to reduce cleaved caspase-3-positive neurons under brain injury (52) and its capacity to ameliorate inflammation at nasal epithelium through intranasal instillation (53) make AG-1478 a suitable candidate to be tested in in vivo AD models. Our study also revealed that the Aurora Kinase A/B/C inhibitor Reversine pretreatment was able to partially restore the survival potential of SH-SY5Y cells upon Aβ treatment. An in-vitro study performed in primary cortical neurons has revealed a potential role of Aurora kinases in neurite initiation, elongation and arborization (54). However, their precise role during AD progression is unknown. Reversine has been proposed as a therapeutic option against cellular senescence (55). This compound increases differentiation potential of progenitor cells toward the neuroectodermal lineage (56). Interestingly, it has been demonstrated that Reversine is able to reactivate insulin pathway, autophagy, glucose uptake and restore glycolytic-Krebs cycle connectivity, rewiring mitochondrial functionality in senescent cells (55). All these processes are considered hallmarks of a neurodegenerative process (57) suggesting Reversine as a potential therapeutic component in AD. Due to AD share common pathologic processes with other neurological syndromes, we cannot rule out the possibility of using AG-1478 and Reversine in therapeutic pipelines against other neurodegenerative diseases. Moreover, due to AD is a progressive neurodegenerative disorder that affects multiple brain regions over time, our omic-based drug repurposing approach is suitable to be applied across brain multiregional post-mortem datasets to evaluate the olfactory dependence of our results. This perspective would help the scientific community to elucidate and classify potential global disease-modifying compounds or region-specific drugs. Follow-up studies are needed to validate and evaluate potential side effects of these candidates in AD patient-derived cells as well as in *in vivo* research.

## Acknowledgments

We are very grateful to the patients and relatives that generously donor the brain tissue for research purposes. We are indebted to the Neurological Tissue Bank from the Hospital Clinic-Institut d’Investigacions Biomédiques August Pi i Sunyer (IDIBAPS, Barcelona, Spain), IDIBELL Biobank and the Biobank from Navarrabiomed for providing us the olfactory specimens as well as the associated clinic-pathological data. Authors thank all JPOST Team for helping with the mass spectrometric data deposit in ProteomeXChange/PRIDE. The Clinical Neuroproteomics Unit of Navarrabiomed is member of the Global Consortium for Chemosensory Research (GCCR) and the Spanish Olfactory Network (ROE) (supported by grant RED2022-134081-T funded by Spanish Ministry of Science and Innovation).

## SUPPLEMENTARY MATERIAL

**Supplementary Table 1.** OT proteomics in AD. The file contains identified and quantified proteins and differential analysis across Braak staging.

**Supplementary Table 2.** Gene Ontology annotations of all differential expressed proteins across Braak stages generated by Metascape

**Supplementary Table 3.** Interlocking of OT differential proteomes with ALZdata repository

**Supplementary Table 4.** Consensual olfactory omics signatures considered in this study.

**Supplementary Table 5.** Reversion potential of olfactory deregulated features in initial AD (by drug-name).

**Supplementary Table 6.** Reversion potential of olfactory deregulated features in initial AD (by drug-type).

**Supplementary Table 7.** Reversion potential of olfactory deregulated features in initial AD (knock-down genes)

**Supplementary Figure 1.** Global OT proteome analysis between non-AD (controls) and AD subjects. Heatmap and Volcano Plots are shown.

**Supplementary Figure 2.** Venn diagram representing the overlap between differential expressed proteins across Braak stages and APP interactors. Differential intensities of APP interactors in each Braak stage is shown. “C” represents intensities from the control group. “AD ini” represents intensities from initial Braak stages (Braak I-II). “AD interm” represents intensities from intermediate AD (Braak III-IV). “AD adv.” represents intensities from advanced AD stages (Braak V-VI stages).

**Supplementary Figure 3.** Venn diagram representing the overlap between differential expressed proteins across Braak stages and Tau interactors. Differential intensities of Tau interactors in each Braak stage is shown. “C” represents intensities from the control group. “AD ini” represents intensities from initial Braak stages (Braak I-II). “AD adv.” represents intensities from advanced AD stages (Braak V-VI stages).

**Supplementary Figures 4-6.** Venn diagram representing the overlap between differential expressed proteins across Braak stages and the proteome previously detected in Aβ plaques. Differential intensities of overlapped proteins in each Braak stage is shown across Supp figure 4, 5 and 6. “C” represents intensities from the control group. “AD ini” represents intensities from initial Braak stages (Braak I-II). “AD interm” represents intensities from intermediate AD (Braak III-IV). “AD adv.” represents intensities from advanced AD stages (Braak V-VI stages).

**Supplementary Figure 7.** Venn diagram representing the overlap between differential expressed proteins across Braak stages and the proteome previously detected in neurofibrillary tangles. Differential intensities of overlapped proteins in each Braak stage is shown. “C” represents intensities from the control group. “AD ini” represents intensities from initial Braak stages (Braak I-II). “AD interm” represents intensities from intermediate AD (Braak III-IV). “AD adv.” represents intensities from advanced AD stages (Braak V-VI stages).

**Supplementary Figure 8.** A) Heatmap representing the functional overlap observed across olfactory areas. B) Circos Plot showing the differential expressed protein overlap across the olfactory bulb (OB), olfactory tract (OT), entorhinal cortex (EC) and hippocampus (HP) in initial AD stages (Braak I-II). Purple lines correspond to differential expressed proteins shared across brain structures.

**Supplementary Figure 9.** Effect of different concentrations of the selected compunds on cell viability of SH-SY5Y cells. Cells were treated with compunds for 24h. Results are presented as % of survival and expressed as mean ± SEM (n ≥ 6). A) SH-SY5Y cell viability upon Indirubin treatment. B) SH-SY5Y cell viability upon GSK3β inhibitor treatment. C) SH-SY5Y cell viability upon AG-1478 treatment. D) SH-SY5Y cell viability upon Reversine treatment.

## AUTHOR CONTRIBUTIONS

Conceptualization, Enrique Santamaría; Data curation, Paz Cartas-Cejudo, Adriana Cortés, Mercedes Lachén-Montes, Elena Anaya-Cubero, Joaquín Fernández-Irigoyen, Enrique Santamaría; Formal analysis: Paz Cartas-Cejudo, Adriana Cortés, Mercedes Lachén-Montes, Elena Anaya-Cubero, Elena Puerta, Maite Solas, Joaquín Fernández-Irigoyen, Enrique Santamaría; Funding acquisition, Joaquín Fernández-Irigoyen, Enrique Santamaría; *In-vitro* studies, Adriana Cortes, Elena Puerta, Maite Solas; Investigation, Paz Cartas-Cejudo, Adriana Cortés, Mercedes Lachén-Montes, Elena Anaya-Cubero, Elena Puerta, Maite Solas, Joaquín Fernández-Irigoyen, Enrique Santamaría; Methodology, Paz Cartas-Cejudo, Adriana Cortes, Mercedes Lachén-Montes, Elena Anaya-Cubero, Joaquín Fernández-Irigoyen, Enrique Santamaría; Writing – original draft, Enrique Santamaría; and all authors gave final approval of the manuscript and are accountable for all aspects of the work.

