## Supplementary material for "Neuropathological stage-dependent proteome mapping of the olfactory tract in Alzheime’s disease: From early olfactory-related omics signatures to computational repurposing of drug candidates": SUPLEMMENTARY FIGURES.pdf

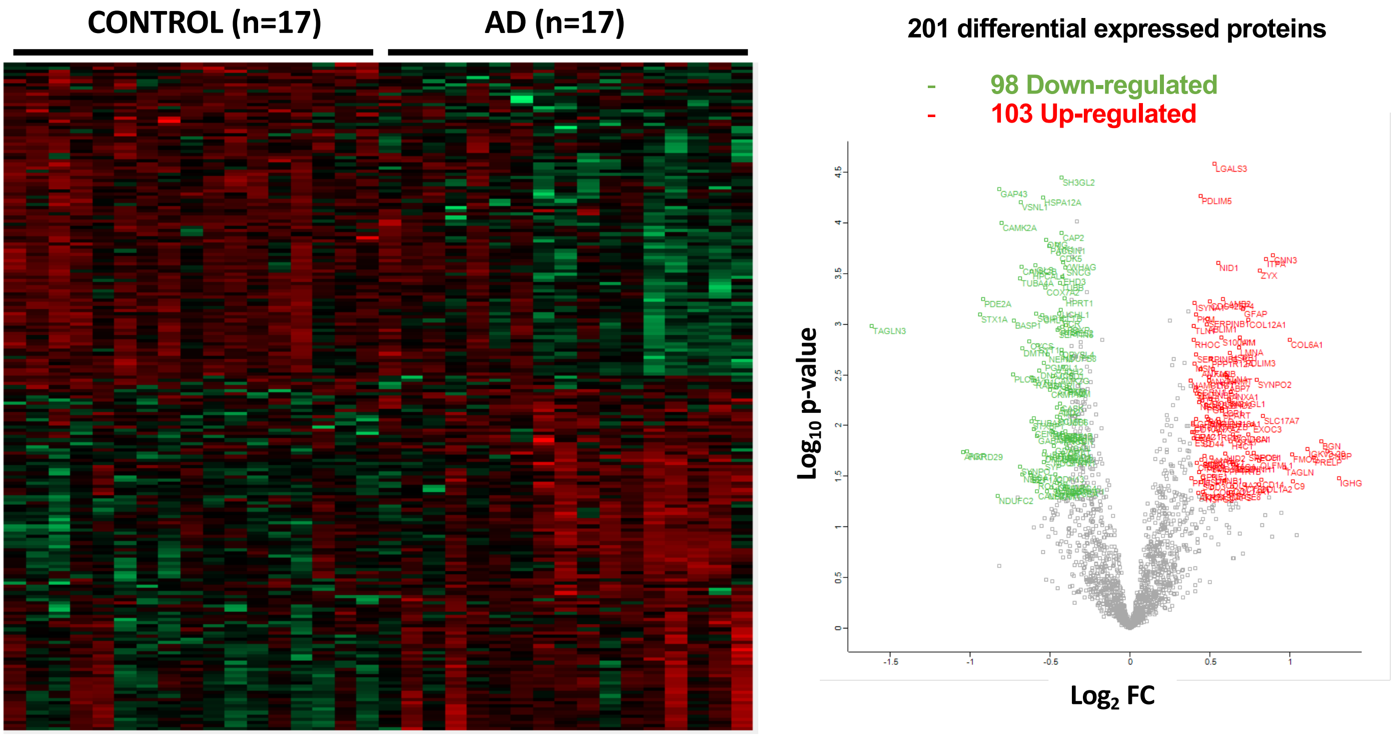


**Supp figure 1**

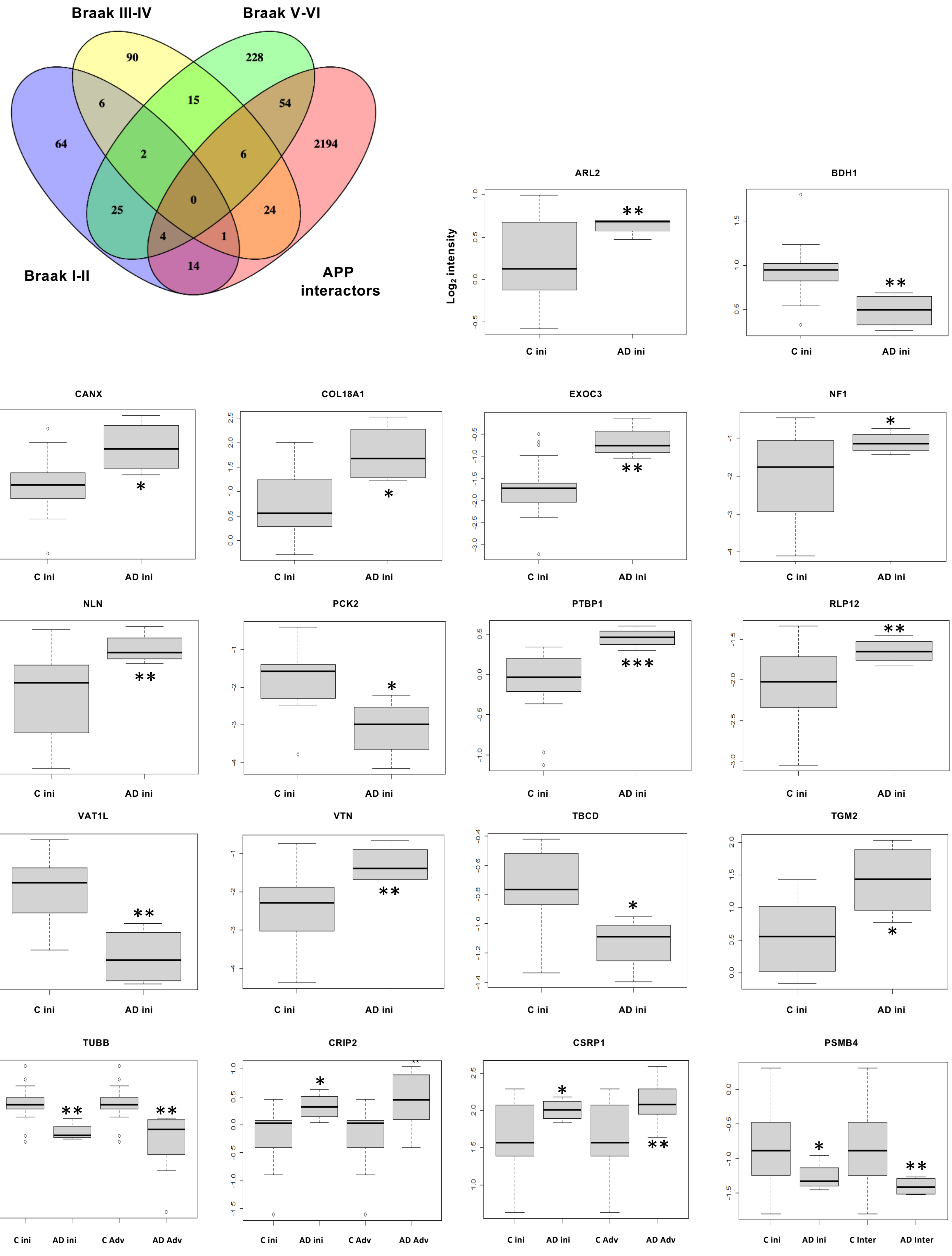


Supp figure 2

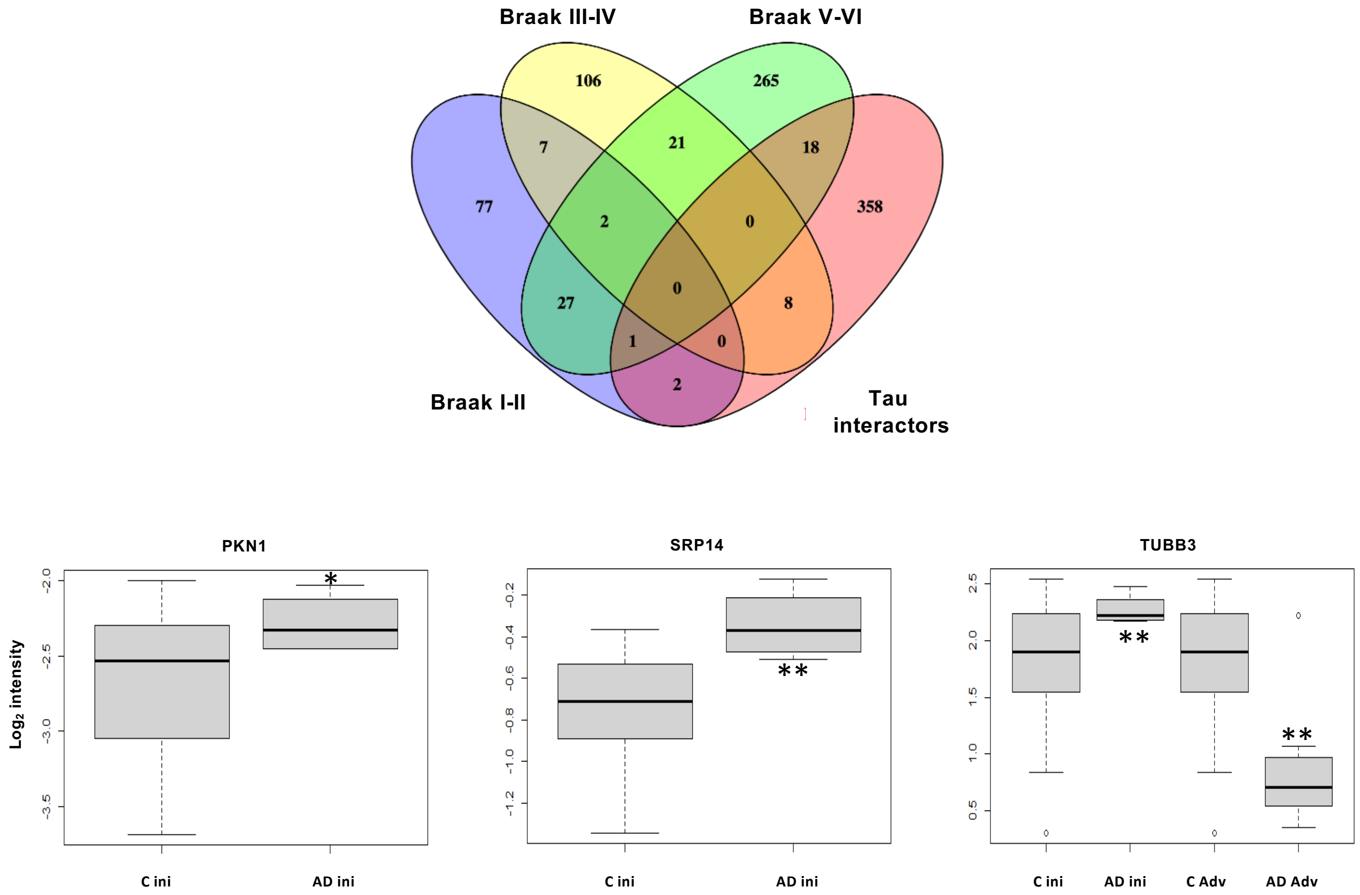


**Supp figure 3**

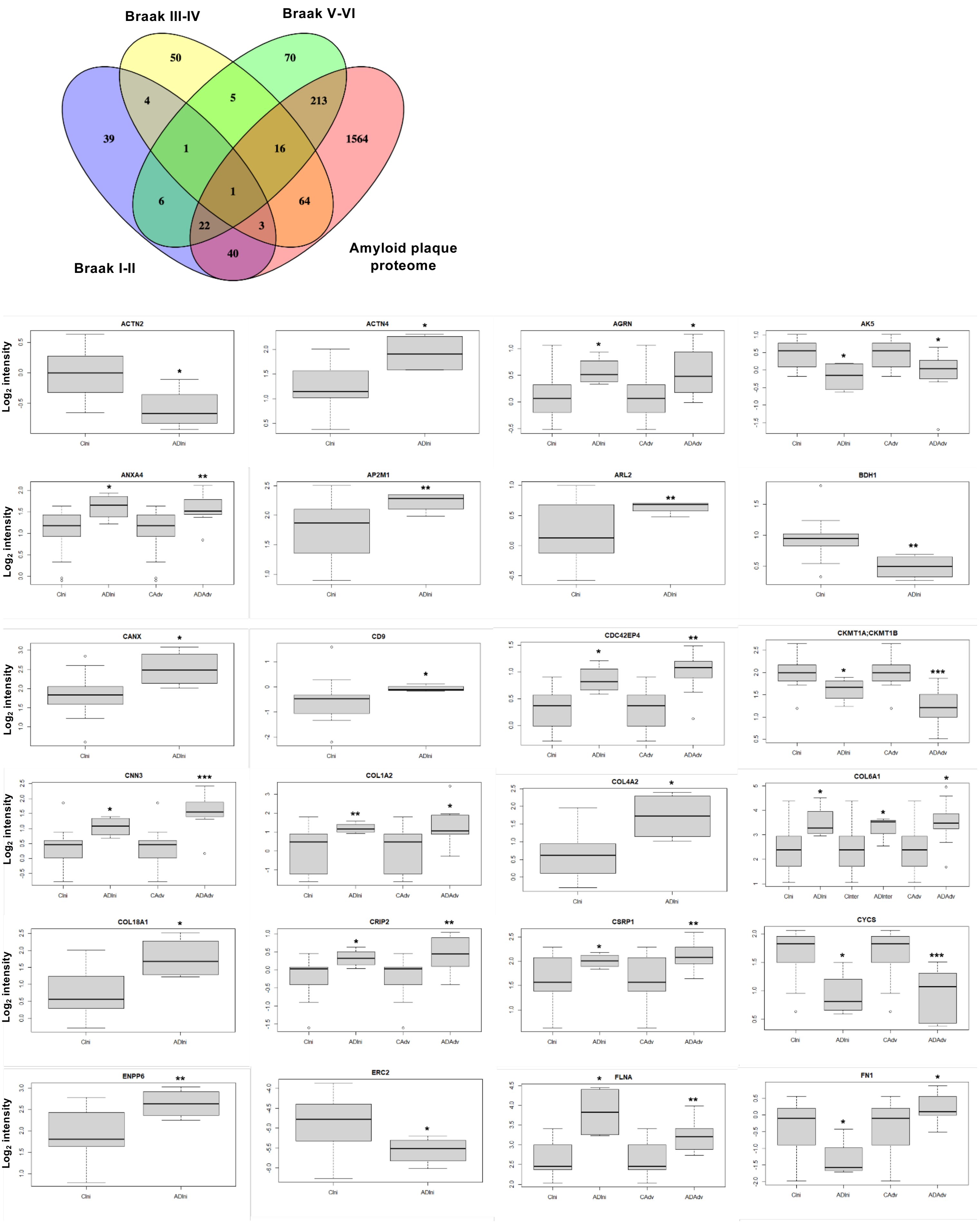


### Supp figure 4

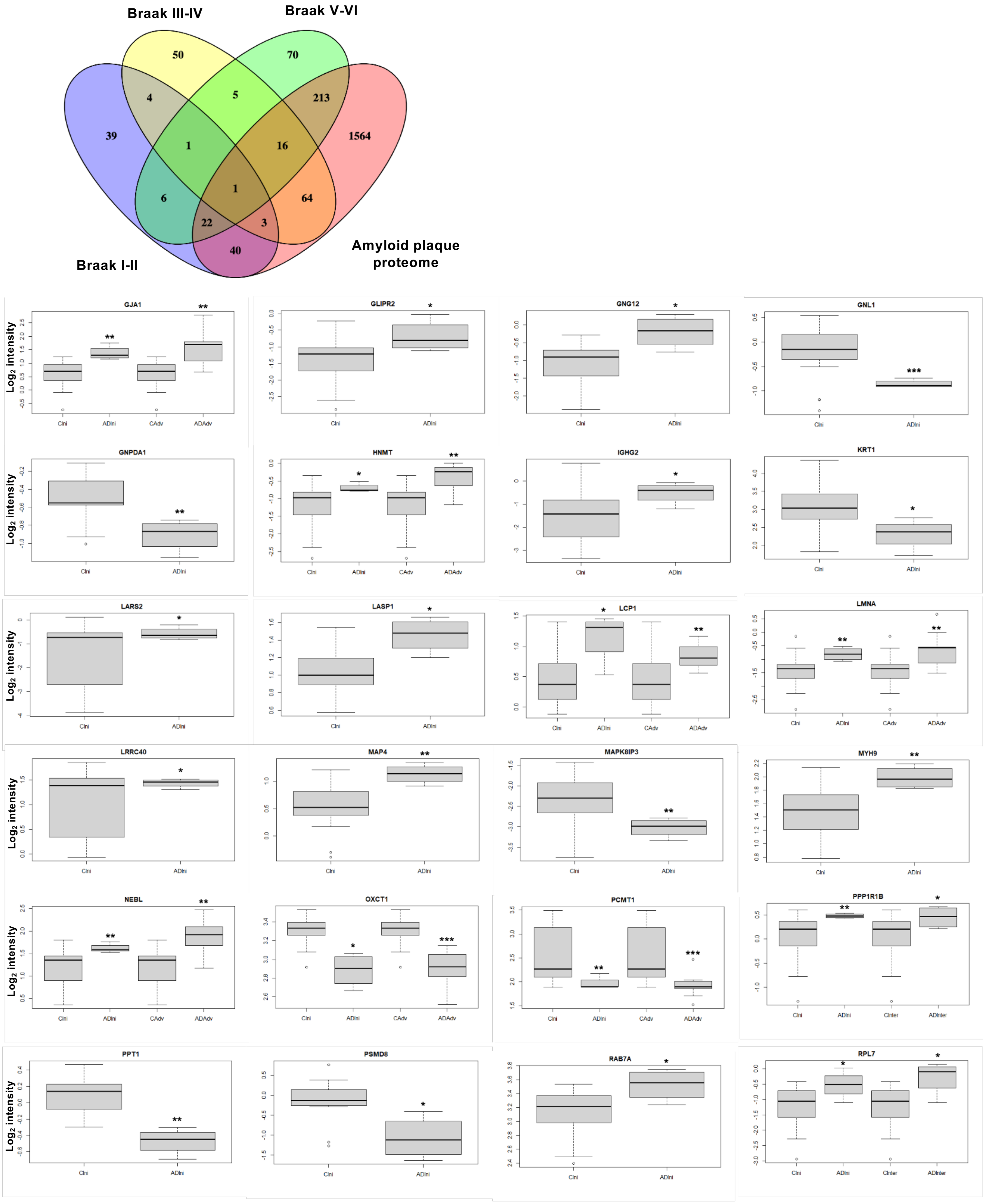


Supp figure 5

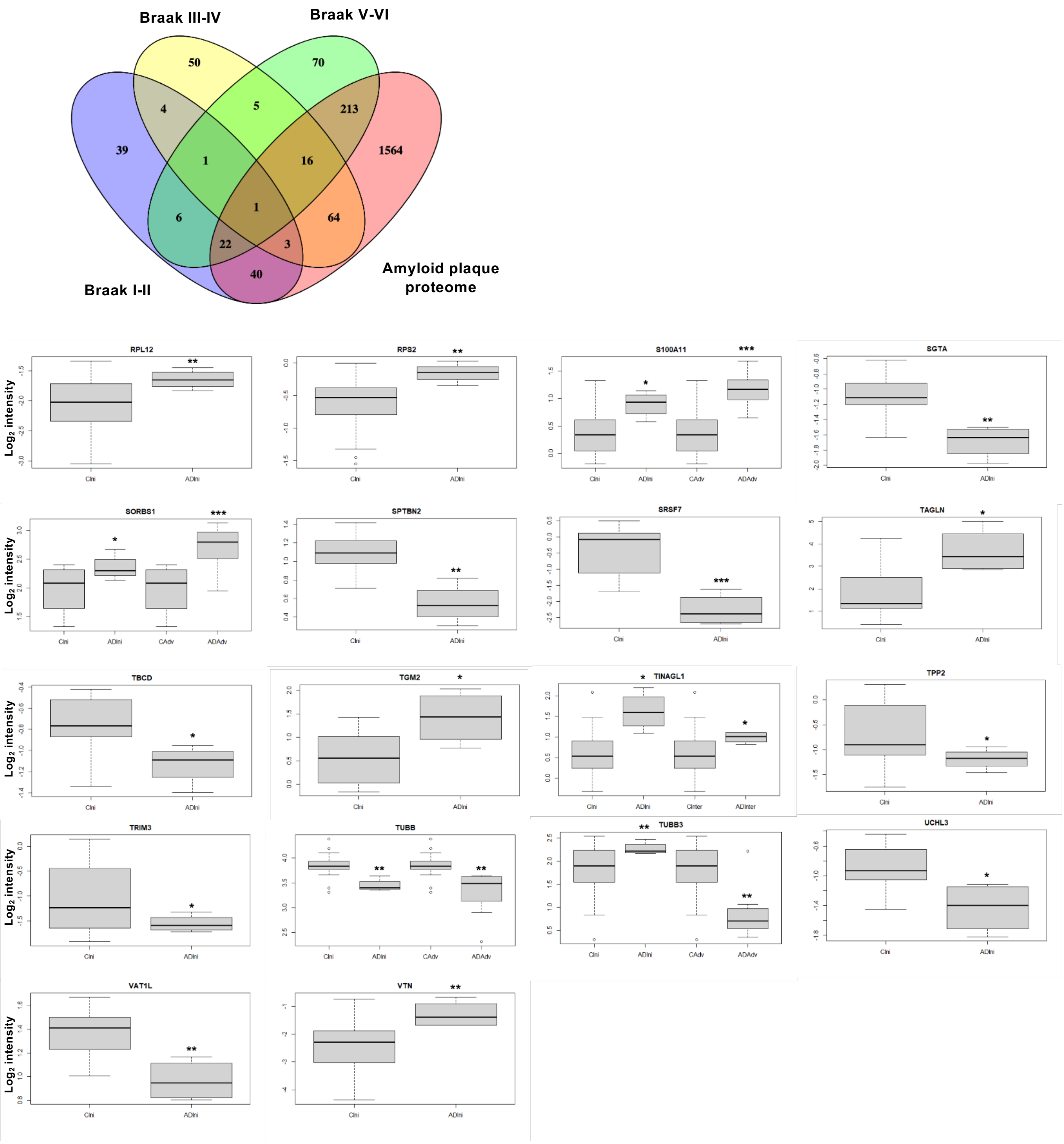


### Supp figure 6

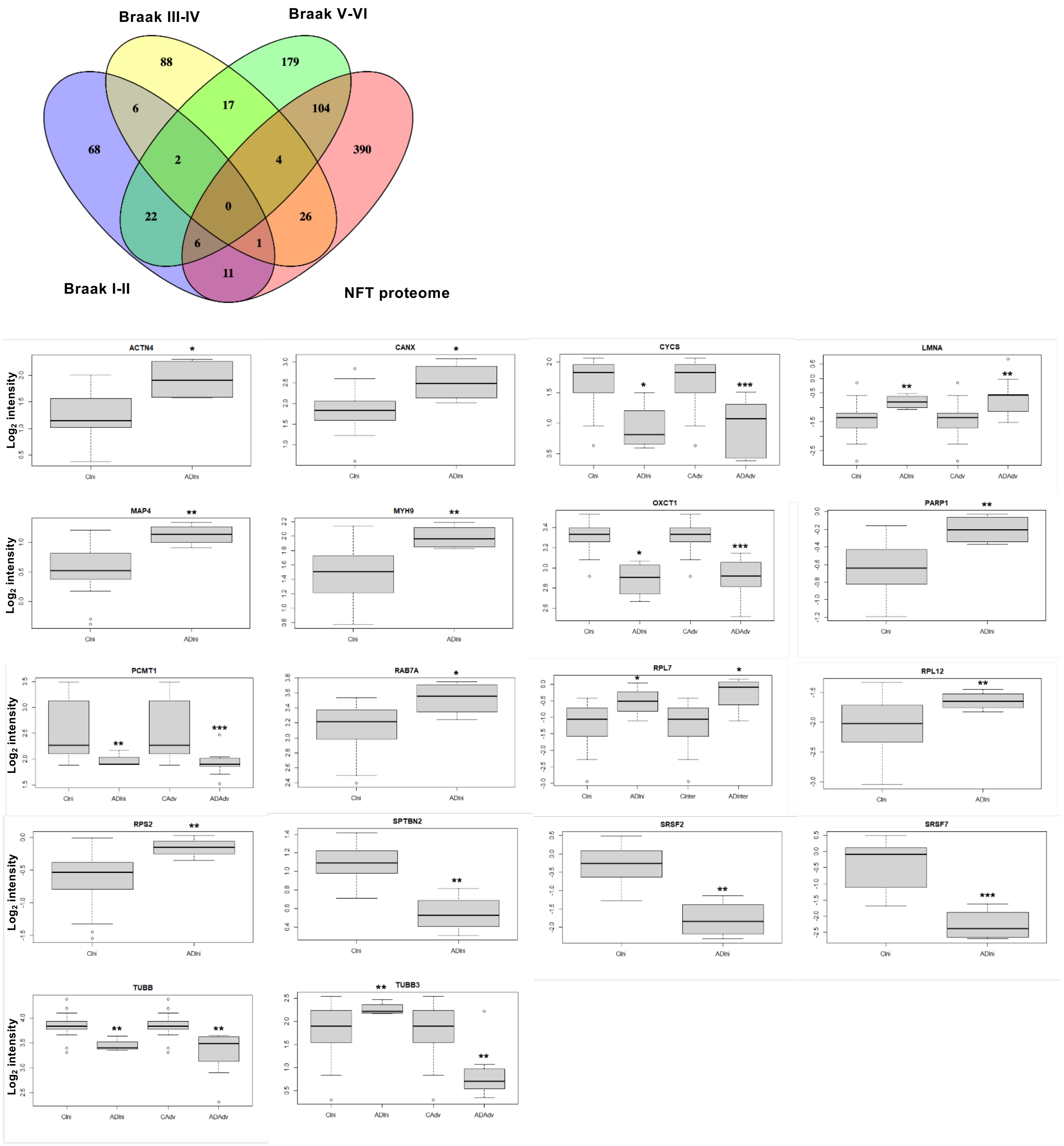


### Supp figure 7

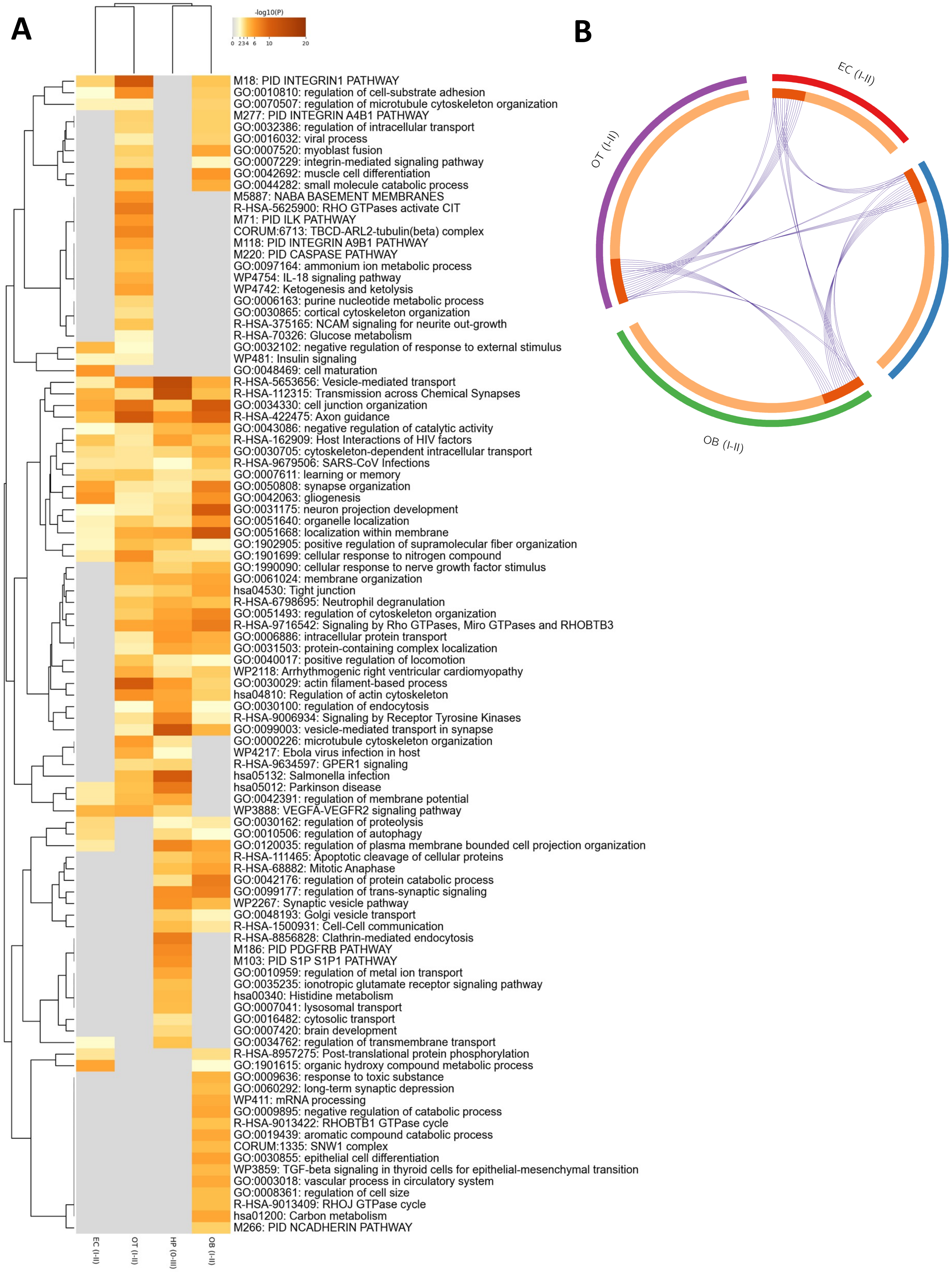


### Supp figure 8

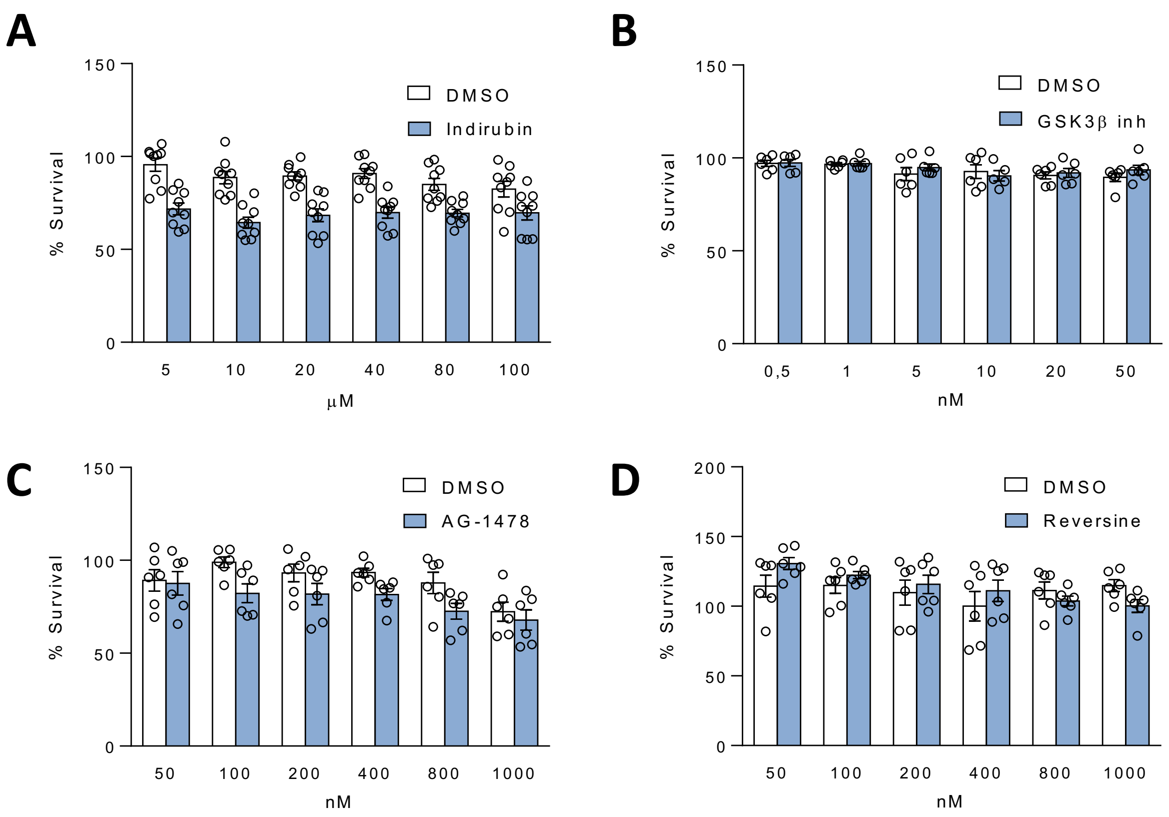


**Supp figure 9**
